# Cell Type-Specific Remodelling of the Rat Hippocampus by Parity and Age

**DOI:** 10.1101/2025.09.17.676894

**Authors:** Andrew J McGovern, Paula Duarte-Guterman, Liisa AM Galea

**Author notes:** Corresponding Author: Liisa AM Galea –.

## Abstract

**Background:** The hippocampus undergoes extensive cellular remodelling throughout life. Parity, the experience of pregnancy and motherhood, triggers profound hormonal changes which modulate brain plasticity in the short and long term. Signatures of past parity are seen in the middle-aged hippocampus in humans and rodents, but how parity shapes cellular composition long-term after pregnancy have not been systematically examined using quantitative, cell-type-specific approaches.

**Methods:** We performed cell type deconvolution on bulk RNA-sequencing data from female rat hippocampus, comparing nulliparous and parous females across age (7 or 13 months either 30 days or 7 months after parturition). We harmonized 349 cell type annotations from three single-cell reference datasets into 27 biologically coherent categories using female-only data. Three-way ANOVA identified independent and interactive effects, while complementary analyses (random forest, PCA, DESeq2) identified parity-associated transcriptional signatures. Cell-specific functional enrichment employed weighted gene set meta-analysis across multiple pathway databases.

**Results:** Age emerged as the dominant factor, significantly altering six cell types, particularly somatostatin (SST) and parvalbumin/Vip interneurons. Regional effects (dorsal and ventral hippocampus) affected nine cell types, while age x region interactions identified two cell types. Parity independently affected three populations: dorsal CA3 pyramidal neurons, SST interneurons, and astrocytes. Cell-type-specific pathway analysis revealed distinct mechanisms including protein degradation in CA3 neurons, stress-response regulation in astrocytes, and disrupted GPCR/signalling-receptor programs in SST interneurons.

**Conclusions:** Our study shows that parity selectively remodels hippocampal cellular architecture through distinct, cell-type-specific molecular programs operating independently of age and region, establishing parity as a critical biological variable in aging research.

## 1. Introduction

The hippocampus exhibits well-established cellular heterogeneity that varies with age and anatomical regions^1^. Distinct subregions of the hippocampus including the dentate gyrus, CA1, and CA3 show distinct cellular compositions across the dorsoventral axis, which are functionally distinct^2^. Age-dependent changes in the hippocampus include increased microglial proportions and altered oligodendrocyte and astrocyte populations^3,4^. However, these studies have been verified predominantly in males, and without dorsoventral investigation.

Pregnancy and motherhood induce profound physiological changes including dramatic hormonal fluctuations, increased metabolic demands, and immune system adaptations^5^. Imaging studies in humans reveal that pregnancy and motherhood result in grey matter volume reductions in the hippocampus that persist for at least 6 years postpartum^6–9^, and further decreased brain aging in middle-age^10,11^. These findings are mirrored in rodent studies which document reduced hippocampal neurogenesis in the short term, followed by increased neurogenesis and synaptic proteins in the hippocampus of middle-aged rats^12,13^. Whether these macroscopic changes reflect uniform tissue remodelling or selective alterations in specific cell populations is unknown. Furthermore, how parity-induced changes relate to established patterns of aging and regional specialization in the hippocampus has not been examined across cell populations. This knowledge gap is significant given psychiatric and neurodegenerative conditions show sex-specific prevalence and onset patterns coinciding with reproductive transitions, and that parity is related to dementia risk^14,15^.

In this study, we ask whether parity is an independent modulator of hippocampal cellular composition, or whether it acts through interactions with aging and hippocampal region. We hypothesized that parity alters cellular composition independently of age and region in the female rat hippocampus. To test this, we performed comprehensive cell-type deconvolution of bulk RNA sequencing from hippocampal samples across parity (nulliparous, biparous, primiparous), age (young (late postpartum), middle-aged), and two hippocampal regions (dorsal, ventral). Cell type deconvolution methods quantify changes in cellular proportions from bulk RNA-sequencing using single-cell reference datasets^16,17^, resolving cell type-specific changes obscured in traditional analyses. However, most brain deconvolution studies use mixed-sex or male-dominated reference datasets, potentially compromising accuracy for female tissues should cell-type specific genes have sex-biased expression. This would distort the basis matrices for deconvolution if male- or mixed- reference sexes are used^18–20^.

Additionally, inconsistent cell type annotations across reference datasets have limited cross- study validation^21^. To address these limitations, we first evaluated whether sex-matched reference datasets improve deconvolution performance for female brain tissue. We then developed a harmonization strategy integrating 349 cell type annotations from three independent single-cell references into 27 biologically coherent categories. Three-way factorial analysis (age × region × parity) revealed powerful effects of age and region as expected. To detect more subtle parity-associated patterns, we employed complementary analytical approaches utilizing differential expression (DESeq2), machine learning classification (random forest), and dimensionality reduction (PCA) to identify genes, and downstream functional pathways, specifically associated with parity. This multi-method strategy enabled us to separate parity effects from the dominant age and regional signals, revealing cell type-specific remodelling unique to parity.

## 2. Methods

### 2.1 Study Design and Data Generation

The experimental design incorporated three critical biological variables: parity, age, and hippocampal region. Thirty female Sprague Dawley rats were obtained from Charles River (Quebec, Canada) and assigned to three parity groups: nulliparous with no reproductive experience, primiparous (one pregnancy and postpartum pup experience), and biparous (two pregnancies and postpartum pup experiences). The pregnancies were timed such that primiparous rats had their first pregnancy when biparous rats had their second pregnancy. For breeding, two females and one male were paired overnight; females were vaginally lavaged each morning and considered pregnant upon confirmation of sperm presence, at which point they were single-housed. Pregnant females were monitored and weighed weekly throughout gestation. One day after birth (postpartum day 1), litters were culled to four males and four females. One female did not have enough pups and was supplemented with pups cross-fostered from other females born on the same day. Original litter sizes and maternal behaviour (time spent licking, nursing, and off-nest, scored between postnatal days 2–8) did not differ significantly between parity or age groups (Supplementary Table S1 in Duarte-Guterman et al. 2023^22^). Rats were euthanized by lethal overdose of sodium pentobarbital between 10:00 and 12:00 at 7 months (30 days postpartum, 7 days after pups had left the nest) or 13 months of age (middle-aged, 7 months after parturition). Brains were rapidly extracted and hippocampi dissected over ice. All protocols were approved by the Institutional Animal Care Committee at the University of British Columbia and conformed to the guidelines set out by the Canadian Council on Animal Care^22^.

### 2.2 RNA isolation, sequencing, and library preparation

Brains were extracted and 300 µm sections were cut on a Leica CM3050S cryostat. Dorsal and ventral hippocampi were punched on dry ice and stored at -80 °C until RNA isolation; sections between -5.2 mm and -6.7 mm Bregma were considered ventral hippocampus^23^. Total RNA was isolated with the Qiagen RNeasy Mini kit per manufacturer’s instructions, resuspended in nuclease-free water, and stored at -80 °C. RNA concentration and purity were determined on a NanoDrop 8000 Spectrophotometer (Thermo Scientific), integrity was verified on a 1% agarose gel and on Agilent Bioanalyzer High-Sensitivity RNA chips, and quantity was confirmed on a Qubit fluorometer. Total RNA was normalised to 500 ng input for the Illumina TruSeq mRNA Stranded library preparation kit. Libraries were sequenced 2 × 75 bp paired-end on an Illumina NextSeq 550 (High Output v2.5 flow cell) at the University of British Columbia Sequencing and Bioinformatics Consortium, targeting ∼25 million reads per sample. The young and aged groups were sequenced in two separate batches. Reads were demultiplexed with Illumina BCL2FASTQ (v2.20.0.422). Adapter sequences and 3′ bases below Q20 were trimmed with Cutadapt and reads shorter than 25 bp were discarded; sequencing quality was confirmed with FastQC (v0.11.8). Trimmed reads were aligned to the rat genome (*\*Rnor_6.0\**, Ensembl release 84) with STAR (v2.6.1a_08- 27), and per-gene read counts were obtained with HTSeq-count (v0.11.0) against the Ensembl 84 GTF. The normalized count matrix was filtered to retain genes meeting mean expression ≥ 1, variance > 0, and detection in ≥ 10% of samples, yielding 12,516 genes for downstream analysis. Raw FASTQ files and per-sample HTSeq read-count tables are deposited at NCBI GEO under accession GSE329776.

### 2.3 Reference Dataset Analysis and Optimization

Three independent mouse single-cell RNA-sequencing reference datasets were used for cross-species deconvolution: the Allen Institute mouse whole-brain 10X Genomics dataset^24^, the mouse-brain Smart-seq2 dataset (NeMO: dat-iye7gkp), and the Yao hippocampus-specific 10X dataset^25^. Mouse references were used for deconvolution of rat bulk RNA-sequencing data, leveraging the high conservation of cell-type-specific gene expression patterns across rodent species and the established practice of cross-species deconvolution.

Each reference was preprocessed to retain hippocampal cells based on anatomical annotations and stratified into female-only, male-only, and mixed-sex configurations. The 10X Genomics dataset contained 22,307 female cells, 59,108 male cells, and 81,964 mixed-sex cells; the Yao hippocampus dataset contained 26,314 / 62,272 / 88,674 cells (female / male / mixed); and the Smart-seq2 dataset contained 3,693 / 1,849 / 5,869 cells. Performance was evaluated using four complementary metrics: Shannon entropy per 1,000 reference cells, calculated as - Σ(pᵢ × log₂(pᵢ))/1,000 where pᵢ is the proportion of cell type *\*i\**; the relative cell-type detection rate; a modified Simpson’s diversity index normalized by log₁₀(n_ref_cells + 1); and detection efficiency (cell types per 1,000 reference cells). On the 10X-based references, female-matched configurations consistently outperformed male-only and mixed-sex configurations: the 10X Genomics female-only reference achieved an entropy of 0.136 per 1,000 cells versus 0.037 (mixed) and 0.048 (male), and the Yao female-only reference achieved 0.096 versus 0.030 (mixed) and 0.036 (male). The Smart-seq2 dataset showed a divergent pattern with the male-only configuration achieving the highest entropy (1.08 vs. 0.41 female and 0.26 mixed), potentially reflecting technical differences in the earlier sequencing platform or male-biased cell-type representation in the original sampling. Based on the consistent female-matched advantage in the larger and more recent 10X datasets, all subsequent deconvolution analyses used female-only reference configurations (reference cell counts and metric definitions tabulated in Supplementary Methods S2.3).

### 2.4 Cell Type Harmonization

The three reference datasets collectively contained 349 cell-type annotations (169 in mouse10x_2020, 125 in mouse_smartseq_2019, and 55 in yao_hippo_10x) with inconsistent nomenclature. To enable cross-dataset comparison, we harmonized them using unsupervised hierarchical clustering of cell-type-specific gene-expression signatures derived from SCDC basis matrices. Pairwise Pearson correlations were computed across the 349 signatures and hierarchical clustering with average linkage was applied to the distance matrix (1 - r) at a height cutoff of 0.2. Harmonization reduced the per-reference annotation count to 16, 10, and 1 categories for mouse10x_2020, mouse_smartseq_2019, and yao_hippo_10x respectively.

Within- and between-cluster separation was confirmed by Wilcoxon rank-sum test (p < 0.001). The full mapping of original-to-harmonized assignments and per-cluster statistics is provided in Supplementary Methods S2.4 and the project repository.

### 2.5 SCDC Deconvolution Analysis

Cell-type proportions were estimated with the Semi-supervised Cell-type Deconvolution with Consistency (SCDC) algorithm, which leverages multiple references simultaneously and provides built-in cross-dataset validation. Animal ID was used as the subject parameter to account for biological replicates; references were ensemble-weighted equally. Sensitivity analyses with alternative weighting schemes produced concordant proportions, and the deconvolution was applied separately to harmonized and non-harmonized cell-type annotations. Convergence checks, reconstruction-error assessments, and replicate-correlation diagnostics are reported in Supplementary Methods S2.5.

### 2.6 Statistical Analysis of Cell Type Proportions

Deconvolved proportions were analysed with a three-way factorial ANOVA model ‘proportion ∼ Age × Region × Parity’. The model tested all main effects, two-way interactions, and the three-way interaction, separately for each of the 23 biologically aligned harmonized cell types, and for individual cell types within each reference with greater than 20 observations or variance greater than 1x10^-8^. Multiple-testing correction used the Benjamini-Hochberg procedure (FDR < 0.05). High-confidence findings were defined by consistent direction and p < 0.05 across at least two reference datasets, and cross-dataset meta-analysis used Fisher’s combined probability test with adjustment for between-dataset correlation (full statistics in Supplementary Methods S2.6).

### 2.7 Differential Gene Expression Analysis

A three-way ANOVA was applied to each of the 12,516 filtered genes (Type III sums of squares, Benjamini-Hochberg FDR < 0.05). To increase power for parity-associated effects, parity was additionally collapsed to a binary factor (nulliparous, *\*n\** = 20, vs. parous, *\*n\** = 40) and tested with DESeq2 (design ‘∼ Age + Parity_Binary’); genes with adjusted p < 0.05 or |fold-change| > 0.3 were retained. Two orthogonal complementary approaches were applied: (i) PCA on the variance-stabilized expression matrix tested the top 10 components (>60% of variance) for factor associations using nested linear models, with PCs significantly associated with parity yielding driver-gene sets (PC5 = 63 genes, PC6 = 128, PC8 = 168); and (ii) a multi-class random-forest classifier (1,000 trees, parity as a 3-level response) ranked parity-predictive genes by Mean Decrease in Accuracy and Mean Decrease in Gini, retaining the top 23 genes by MDA. The full weighted-score formulation, model diagnostics, and feature-selection details are in Supplementary Methods S2.7.

### 2.8 Cell-Specific Functional Enrichment Meta-Analysis

For parity-responsive cell types, weighted gene set enrichment analysis (clusterProfiler v4.8.0, ‘gseGÒ, ‘gseKEGG’, ‘gsePathway’, and ‘GSEÀ functions) was performed across seven pathway database collections: GO:BP, GO:MF, GO:CC, KEGG, Reactome, MSigDB Hallmark, and MSigDB Curated. Four complementary weighting schemes (association- weighted, expression-weighted, statistical-weighted, and composite-weighted) combined gene–cell-type association scores with fold-change magnitude and statistical significance to capture different aspects of pathway response (formulae in Supplementary Methods S2.8).

For each cell type, weighting scheme, and percentage cutoff (top 5% / 10% / 25% by signed weighted score), the ranked gene list was submitted to permutation-based GSEA against each database. Pathway-level magnitude is reported as the normalised enrichment score (NES) from GSEA. Positive NES indicates enrichment in the up-ranked leading edge; negative NES indicates enrichment in the down-ranked leading edge. Input gene lists are derived from the highest-ranked parity-associated genes by signed weighted score (positive scores reflect parity-upregulated genes), pathways with positive NES are interpreted as upregulated in the relevant cell type, and pathways with strongly negative NES as downregulated. Results were integrated through a meta-analysis that scored each pathway by consensus across methods (proportion of methods returning the pathway), evidence strength (weighted mean - log10(FDR)), and NES consistency (1 - coefficient of variation across methods, capped at 0). The composite meta-score = consensus × evidence × consistency, with confidence tiers defined as Ultra-High (≥ 4 methods, meta-score > 7), High (≥ 3 methods, meta-score > 5), Moderate (≥ 2 methods, meta-score > 3), and Method-specific (single-method enrichment). Cell-type-specific pathways were defined as those enriched in ≤ 2 cell types.

### 2.9 Data Integration and Visualization

All analyses were performed in R 4.3.0 with packages SCDC (v0.0.0.9000), DESeq2 (v1.40.0), clusterProfiler (v4.8.0), randomForest (v4.7-1.1), and tidyverse (v2.0.0); visualizations used ggplot2 (v3.4.2). Full software versions, driver scripts are provided in Supplementary Methods (Software and reproducibility). All generated tables, results, and figure generation scripts are present in the project repository at https://github.com/AJMcGovernLab/ReproductiveExperienceAndAgeDeconvolution.

## 3. Results

### 3.1 Sex-Matched Reference Datasets Optimize Cell Type Deconvolution Performance

To optimize parameters for cell type deconvolution, we first evaluated the impact of sex-specific reference dataset composition on deconvolution performance across three independent mouse single-cell RNA-seq datasets: 2019 Smart-seq2, 2020 10x Genomics, and Yao Hippocampus 10x (Figure 1). Female-matched reference datasets demonstrated superior performance across multiple metrics when deconvolving female hippocampal samples. The 10x Genomics female-only reference (n=22,307 cells) achieved higher entropy per 1,000 reference cells (0.136) compared to mixed-sex (0.037, n=81,964 cells) or male-only (0.048, n=59,108 cells) configurations (Figure 1b). The female-only reference detected 67 cell types compared to 95 in mixed and 73 in male-only references but showed superior detection efficiency with 3.00 cell types per 1,000 reference cells versus 1.16 (mixed) and 1.24 (male) (Figure 1d). The Yao Hippocampus dataset showed similar patterns. Female-only reference (n=26,314 cells) detected 1.94 cell types per 1,000 cells compared to 0.62 (mixed, n=88,674 cells) and 0.77 (male-only, n=62,272 cells). Female-only reference achieved entropy per 1,000 cells of 0.096 versus 0.030 (mixed) and 0.036 (male). The Smart-seq2 dataset showed different patterns, with male-specific reference achieving higher entropy per 1,000 cells (1.08, n=1,849 cells) compared to female-only (0.41, n=3,693 cells) and mixed (0.26, n=5,869 cells), potentially reflecting technical differences between 10x genomics and Smart-seq, or male biases in this earlier sequencing technology. Based on these performance metrics showing consistent female advantage in 10x datasets, we selected female-only reference datasets for all subsequent deconvolution analyses.

**Figure 1.**
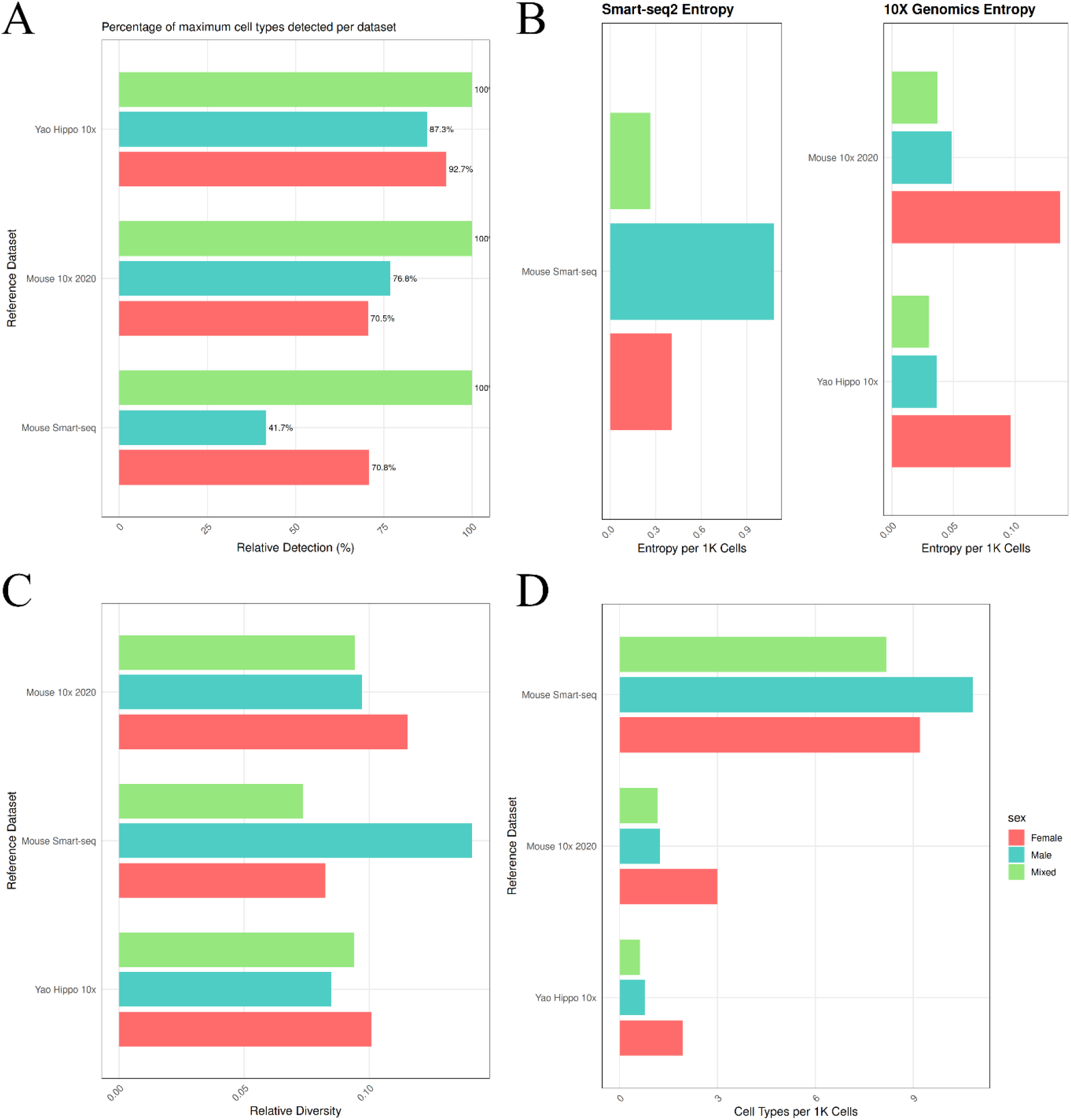
Sex-Matched Reference Datasets Outperform Mixed. Multiple scRNAseq reference datasets (2019 Smart-seq, 2020 10x, Yao Hippo) were tested to determine if sex was an influencing factor in cell type deconvolution. Each dataset was first filtered to ensure it only included cells located in the hippocampus to match our study. (A) Each dataset was used to deconvolute the bulkRNAseq in mixed, male-only, and female- only compositions. Mixed reference datasets always produced more detected cell types as a natural consequence of differences in cell types available in male and female data. (B) Female-matched reference data deconvoluted more cell type Shannon entropy (a measure of cell type diversity) relative to the number of reference cells from the 10x genomics, but male-only reference data outperformed mixed and female-matched from the smart-seq reference data. (C) Similarly, female-matched 10x genomic reference data produced more relative diversity than male-specific or mixed reference data sets. (D) Notably, mixed and female-matched reference data outperformed male-specific data in the YaoHippo 10x dataset. The 2019 Smartseq reference dataset produced much greater diversity from the male-specific reference data than either female-matched or mixed. The number of cell types detected per 1000 reference cells revealed female-matched greatly outperformed mixed or male- specific reference data from the 10x genomics datasets. However, this may be an artifact of a finite number of cells to detect and a lower number of female reference cells. The smart-seq reference dataset detected more cell types from male-specific datasets.

### 3.2 Hierarchical Clustering Harmonizes 349 Cell Types into 27 Biologically Coherent Categories

The three reference datasets collectively contained 349 distinct cell type annotations with inconsistent nomenclature across sources. To enable meaningful cross-dataset comparisons and biological interpretation, we developed a harmonization strategy using unsupervised hierarchical clustering of cell type gene expression signatures (Figure 2). Pairwise correlation analysis of cell type-specific gene expression profiles revealed strong clustering patterns corresponding to major cell lineages. The correlation matrix showed clear block structures with mean within-cluster correlations of r = 0.745 ± 0.157 versus between-cluster correlations of r = 0.634 ± 0.153 (Wilcoxon rank-sum test p < 0.001), validating the biological coherence of identified clusters. Hierarchical clustering with a height cutoff of 0.2 produced 27 dataset-specific harmonized categories across the three references (16 in mouse10x_2020, 10 in mouse_smartseq_2019, and 1 in yao_hippo_10x), corresponding to 23 unique biological categories when deduplicated across datasets. UMAP visualization confirmed that unsupervised clustering preserved major cell class organization, with neuronal populations forming distinct clusters separate from glial populations, vascular cells, and immune cells (Figure 2d-e).

**Figure 2.**
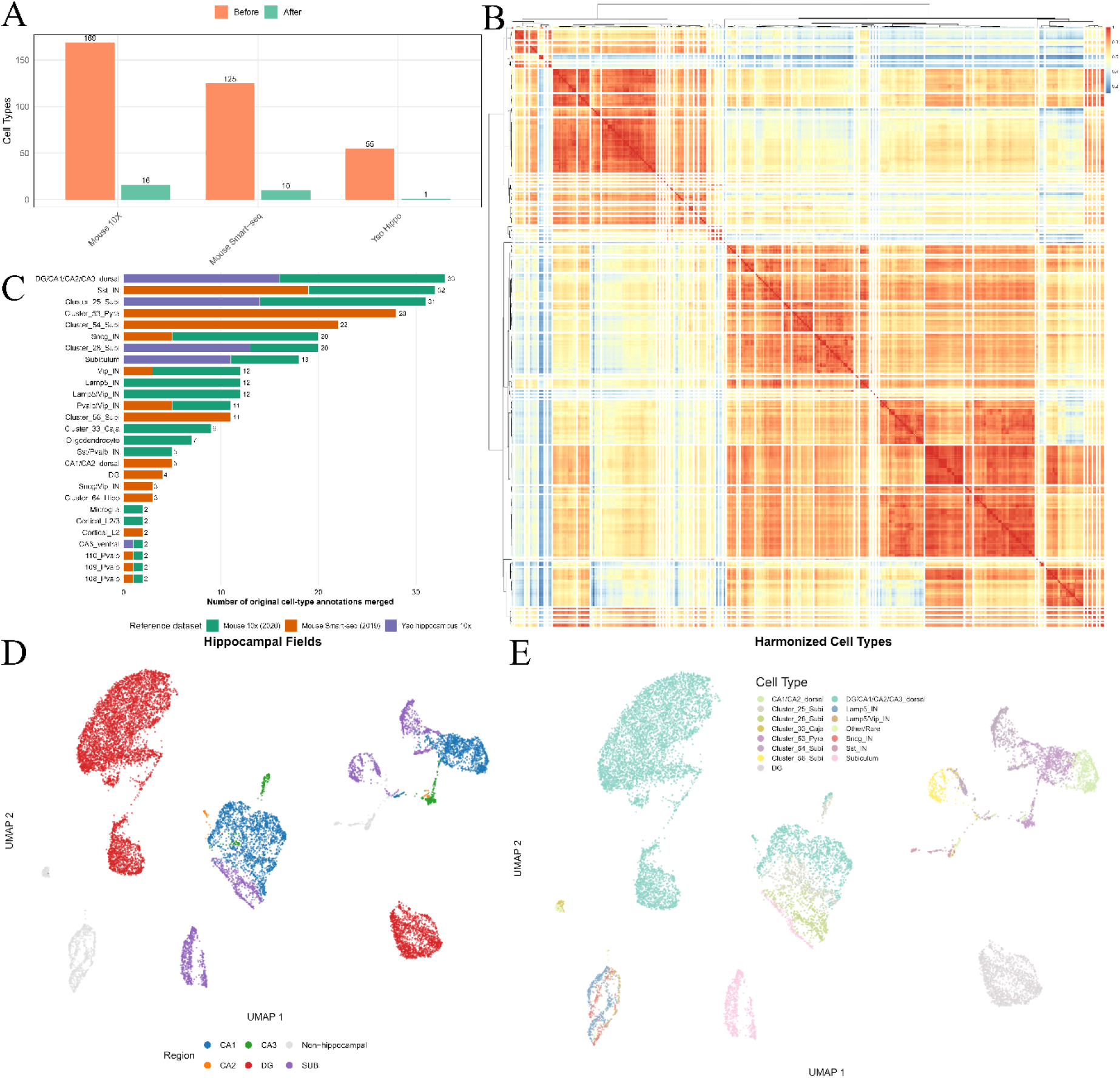
Cell Type Harmonization Across Reference Datasets. (A) We reduced our 349 cell types across three reference datasets down to 27 cell-types to improve the interpretability of our deconvolution analysis using (B) hierarchical clustering based on gene expression. The clusters were then named based on the most common naming patterns within each cluster. (C) This resulted in clusters containing varying numbers of cell types such as the largest cluster DG/CA1/CA2/CA3_dorsal, and second largest Sst_IN, which merged constituent cell types from across datasets. UMAPs show that organization based on cell class definition is largely maintained by unsupervised hierarchical clustering by region (D) and cell types (E).

### 3.3 Age Dominates Hippocampal Cellular Remodelling with Region-Specific Interactions

#### 3.3.1 Deconvolution

Three-way ANOVA of harmonized cell type proportions (age × region × parity) identified significant effects on hippocampal cellular composition (Figure 3a, Supplementary Figure 1). Six cell types showed significant age-related changes (Figure 3b): Sst_IN (p = 3.66 × 10⁻⁸; - 22.4% with age), 386_Micro-PVM (p = 7.30 × 10⁻¹⁰; -38.0%), Pvalb/Vip_IN (p = 0.0029; - 91.4%), Microglia (p = 0.0066; only found in aged samples), Subiculum (p = 0.015; -24.8%), and Cluster_28_Subi (p = 0.021; +16.7%). Nine cell types showed significant dorsal-ventral differences (Figure 3c): DG/CA1/CA2/CA3_dorsal (p = 6.51 × 10⁻²⁹; +16.1% dorsal), Sst_IN (p = 4.14 × 10⁻¹⁵; -32.6% ventral), Oligodendrocyte (p = 4.95 × 10⁻¹⁶; +45.3% dorsal), Cluster_25_Subi (p = 9.32 × 10⁻¹⁹; -39.9% ventral), Subiculum (p = 2.57 × 10⁻⁹; -54.4% ventral), CA1/CA2_dorsal (p = 4.69 × 10⁻⁴; dorsal), 386_Micro-PVM (p = 3.79 × 10⁻⁴; - 21.0% ventral), Pvalb/Vip_IN (p = 0.0087; -84.5% ventral), and Sncg_IN (p = 0.018; -17.3% ventral). Two cell types showed significant age × region interactions: 386_Micro-PVM (p = 0.003) and Oligodendrocyte (p = 0.013), indicating region-specific aging patterns (Figure 3d).

**Figure 3.**
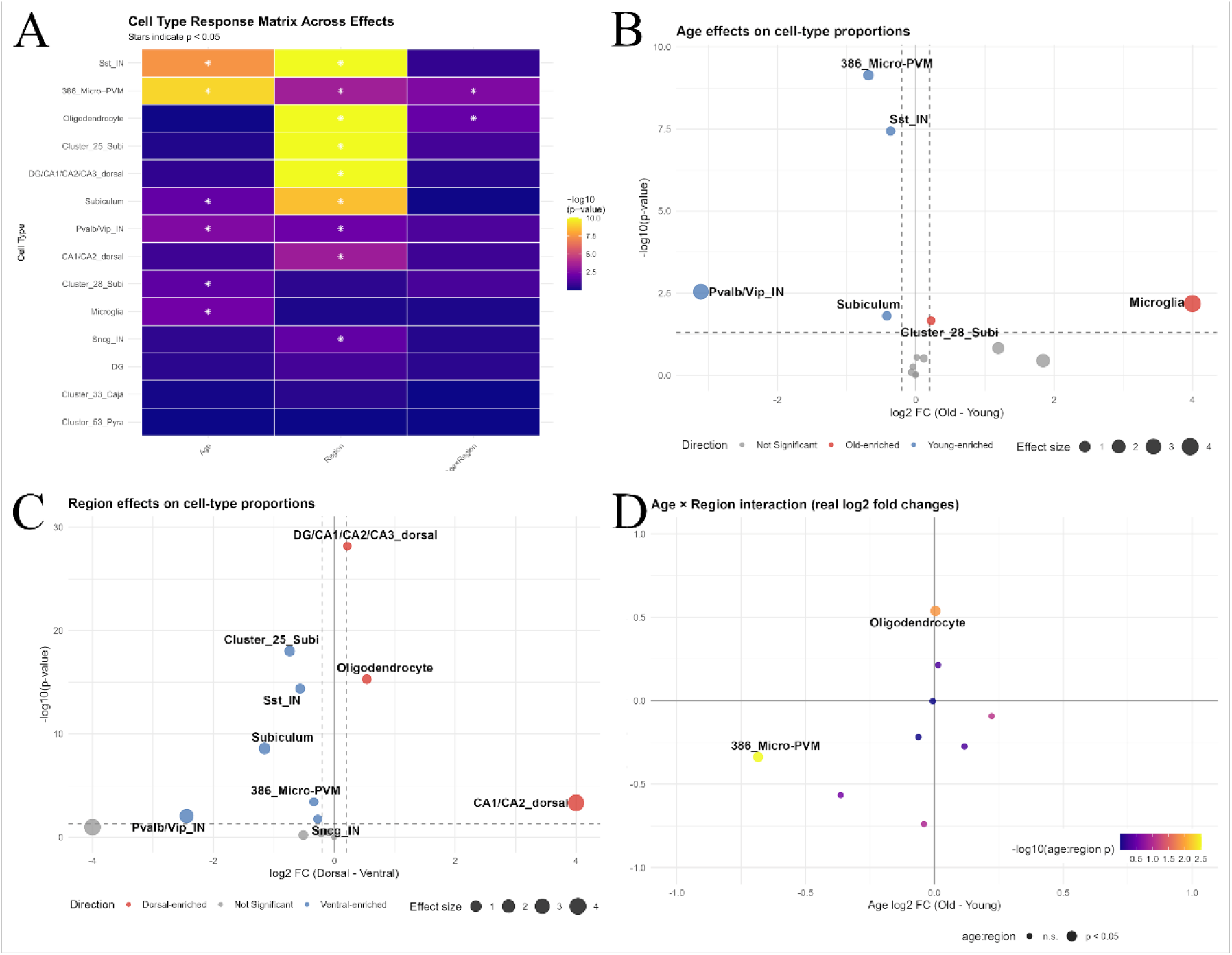
Age, Region, And Age-Region Interactions Regulate Cell Proportions in the Hippocampus. (A) A three-way ANOVA of cellular proportions identified significant effects of age, region, and age × region interactions on cell-type composition in the female hippocampus. (B) Age altered the abundance of six cell types: Sst_IN, Pvalb/Vip_IN, 386_Micro-PVM, and Subiculum showed lower proportions in aged hippocampi, Cluster_28_Subi showed higher proportions in aged hippocampi, and Microglia showed an age effect at p = 0.0066 with directionality unstable due to near-zero detection in young samples. (C) Nine harmonized cell types were significantly influenced by regional differences: DG/CA1/CA2/CA3_dorsal, Oligodendrocyte, and CA1/CA2_dorsal showed greater abundance in the dorsal hippocampus, whereas Sst_IN, Pvalb/Vip_IN, Sncg_IN, Subiculum, Cluster_25_Subi, and 386_Micro-PVM showed greater abundance in the ventral hippocampus. (D) Age × region interactions identified two cell types (386_Micro-PVM and Oligodendrocyte) with region-specific aging trajectories.

#### 3.3.2 Functional Enrichment Analysis of Age and Region Associated Deconvoluted Cell Types

A three-way ANOVA (age × region × parity) of 12,516 genes homologous to mice and rats identified transcriptional regulation with regional effects dominating the transcriptomic landscape. Age effects were extensive, significantly altering 8,163 genes (65.2% of genome, FDR < 0.05) (SuppFig 1a). Region emerged with main effects significantly altering 6,086 genes (48.6% of transcriptome, FDR < 0.05) (SuppFig 1b), while only 8 genes were identified with significant parity main effects (FDR < 0.05) (SuppFig 1c). Age-region interactions affected 1,798 genes (14.4%) (SuppFig 1d). Higher-order interactions involving parity were absent with only 7 genes showing age × parity interactions at FDR<0.05, and none reached significance for parity × region or three-way interactions. Effect overlap analysis showed that 5,268 genes displayed region-only effects, 980 genes showed age:region-only interactions, and 818 genes exhibited both effects, totalling 7,066 unique differentially expressed genes.

Weighted gene set enrichment analysis was performed using four complementary methods (association-weighted, expression-weighted, statistical-weighted, and composite-weighted) across seven pathway databases. Meta-analysis identified high-confidence pathways supported by multiple methods, with confidence tiers based on method agreement and statistical significance (Figure 4a). A cell type vulnerability hierarchy was identified: Sst_IN (age) > Pvalb/Vip_IN (age) > 386_Micro-PVM (age-region) > CA1/CA2_dorsal (region) > Oligodendrocyte (age-region), based on mean meta-scores and high-confidence pathway counts. Age effects showed the strongest enrichment signatures, with Sst_IN displaying the highest vulnerability score (8.19) and 31 high-confidence pathways (Figure 4b-c). Ultra-high confidence pathways (meta-score >7, all 4 methods) in Sst_IN included catalytic activity (meta-score = 7.11, meta-FDR = 2.69 × 10⁻¹⁹). High-confidence pathways included ion binding (meta-score = 5.27), catalytic activity acting on protein (meta-score = 5.22), and ATP-dependent activity (meta-score = 5.15). Pvalb/Vip_IN showed more limited age effects with 5 high-confidence pathways, including ATP-dependent activity (meta-score = 3.43) and peptide ligand-binding receptors (meta-score = 3.07) (Figure 4d). The 386_Micro-PVM population showed one high-confidence pathway: complex of collagen trimers (meta-score = 3.01) (Figure 4e). Region effects were most pronounced in CA1/CA2_dorsal neurons, with 3 high-confidence pathways related to extracellular matrix organization: external encapsulating structure (meta-score = 3.75), extracellular matrix (meta-score = 3.75), and collagen-containing extracellular matrix (meta-score = 3.61) (Figure 4f). Age-Region interactions showed significant enrichment in two cell types. 386_Micro-PVM exhibited high confidence pathways in signalling receptor activity (meta-score = 7.00) and molecular transducer activity (meta-score = 7.00), with additional moderate-confidence pathways in transmembrane signalling (meta-score = 4.97) and G protein-coupled receptor activity (meta-score = 4.00) (Figure 4g). Oligodendrocytes showed moderate-confidence enrichments in signalling receptor activity (meta-score = 4.47) and neuroactive ligand-receptor interaction (meta-score = 3.22) (Figure 4h).

**Figure 4.**
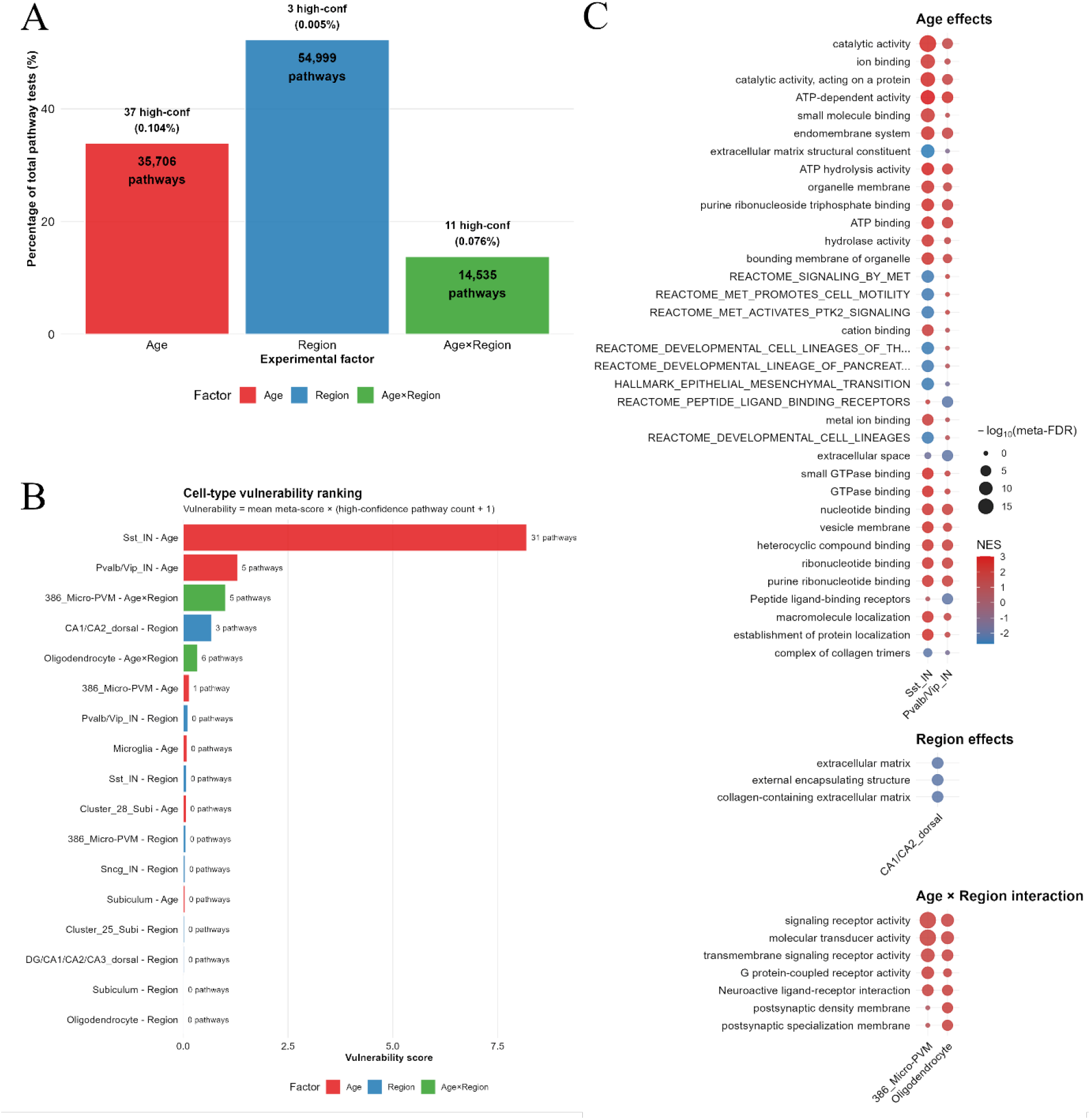
Cell Type Specific Enriched Functional Enrichment. (A) Factor distribution showing the relative contribution of aging, regional, and age×region interaction effects to the enriched pathway landscape. Bar plot displays the percentage of total pathways (n=105,240) attributable to each experimental factor, with annotations indicating the number of pathways and high-confidence enrichments per factor. (B) Cell-type vulnerability ranking across experimental factors. Vulnerability scores calculated as meta-score × (high-confidence pathways + 1), integrating pathway enrichment strength with the number of significantly enriched pathways. Horizontal bars show the top 15 cell type-factor combinations, with annotations indicating the number of high-confidence pathways for each combination. (C) Dot plots showing factor related cell-specific enrichment based on vulnerability score and at least 1 cell-type specific high-confidence pathway. Point size represents the statistical significance of the pathway (−log_10_meta-FDR), with colour coding indicates upregulation (red) or downregulation (blue) in network enrichment score (NES).

#### 3.4 Parity Effects on Specific Cell Populations in the Hippocampus

Parity was not identified to significantly influence any harmonized cell type proportions, gene expression with age- and region- interactions, and revealed only 8 genes to be significantly different between nulliparous, biparous, and primiparous groups. To further explore parity effects, we increased the power of the analysis by combining the biparous and primiparous groups, and tested changes in cell-type proportions across the unharmonized cell-type identities. Enriched genes in parity were identified utilizing DESeq2 with age as a covariate, random forest modelling with elbow analysis, and principal component feature enrichment with downstream gene loading identification.

Comprehensive DESeq2 analysis across the entire hippocampal dataset (design: age + parity_binary) identified genes with nominal statistical significance (p < 0.05) that were subsequently weighted and integrated with PC and random forest results (Supplementary Figure 2) for functional enrichment analysis (Supplementary Figure 3). Analysis of the principal components through binary parity analysis (nulliparous vs parous) identified PC5, PC6, and PC8 as significantly associated with parity through factor association testing. For each parity-associated PC, driver genes were identified by testing genes ranked by absolute loading magnitude for their contribution to the PC-factor association quantified by calculating the reduction in model R² when that gene’s influence was removed from PC scores. This approach yielded PC-specific driver gene sets: PC5 (63 genes), PC6 (128 genes), and PC8 (168 genes; Supplementary Figure 4). Random forest modelling identified the top 23 genes, ranked by MDA with feature importance scores ranging from 3.04 to 1.83 for the most predictive markers, which were selected as the most informative parity-predictive features, with this number chosen based on the elbow point in the importance score distribution (Supplementary Figure 5).

#### 3.4.1 Deconvolution

Analysis of individual datasets identified four cell identities showing significant parity effects (p < 0.05; Figure 5a). These effects were independent of age and regional programs, with no significant three-way interactions detected. Dorsal CA3 pyramidal neurons showed parity-associated increases across datasets: mouse10x_2020 356_CA3-do (p = 0.010, mean proportion = 6.76%, +8.7% in parous) and mouse_smartseq_2019 358_CA3-do (p = 0.0086, mean proportion = 63.7%, +4.4% in parous). The astrocyte populations (376_Astro in mouse10x_2020) showed a parity-associated decrease (p = 0.022, mean proportion = 28.7%, -1.8% in parous). The Sst interneurons (78_Sst HPF in mouse10x_2020) showed a parity-associated increase (p = 0.029, mean proportion = 1.64%, +8.1% in parous) (Figure 5b). Directional log2 fold change estimates were consistent across three independent estimators (estimated marginal means, additive linear-model coefficients, and per-stratum log2 fold change averaging). CA1 hippocampal neurons approached significance for parity effects (p = 0.052). No significant parity × age or parity × region interactions were detected, however Sncg GABAergic interneurons approached parity × age significance (p = 0.064).

**Figure 5.**
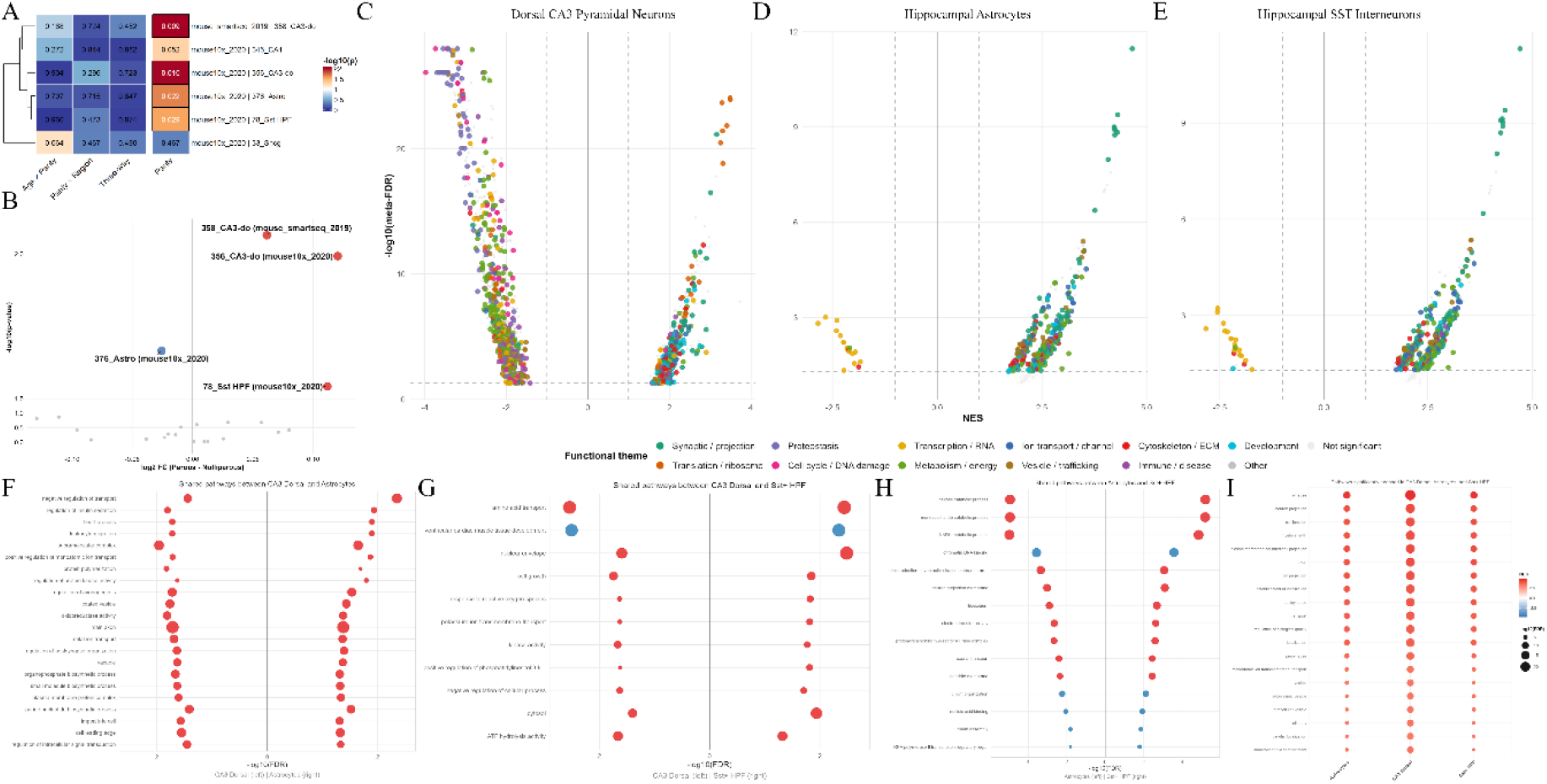
Parity influences dorsal CA3, SST interneuron, and astrocyte proportions through cell-type- specific and shared pathway programs. **(A)** Heatmap of parity-related effects on cell-type proportions. **(B)** Volcano plot of parity effects on cell-type proportions across all deconvolved cell-type × dataset combinations. The four parity-significant populations are coloured by direction (red = higher in parous, blue = higher in nulliparous; grey = not significant) and labelled: dorsal CA3 pyramidal neurons (358_CA3-do, mouse_smartseq_2019; 356_CA3-do, mouse10x_2020) and SST interneurons (78_Sst HPF, mouse10x_2020) increased, whereas astrocytes (376_Astro, mouse10x_2020) decreased in parous animals. **(C-E)** Pathway directional volcano plots for the parity-responsive cell types: dorsal CA3 pyramidal neurons (**C**; 358 Smart-seq reference), hippocampal astrocytes (**D**; 376_Astro), and hippocampal SST interneurons (**E**; 78_Sst HPF). For each pathway the x-axis shows the mean normalised enrichment score (NES) (NES > 0 = upregulated, NES < 0 = downregulated in parous animals); dashed lines mark meta-FDR = 0.05 and |NES| = 1. Points are coloured by functional theme (shared legend). CA3 neurons show downregulated proteostasis and cell-cycle programs alongside upregulated translation/ribosome pathways. Astrocyte pathways are predominantly upregulated, and SST interneurons show both up- and down-regulated programs. **(F–H)** Two-sided dot plots of pathways co- enriched between pairs of parity-responsive cell types: (**F**) CA3-astrocytes, (**G**) CA3-SST interneurons, and (**H**) astrocytes-SST interneurons. Each shared pathway is plotted for both cell types, the first on the left and the second on the right (positive x-axis). Horizontal position encodes -log₁₀(FDR), dot size encodes absolute NES, and colour encodes direction (red = upregulated, blue = downregulated in parous animals). **(I)** Dot plot of the top 20 pathways shared across all three parity-responsive cell types. Colour intensity encodes NES and dot size encodes -log₁₀(FDR) with pathways are ordered by mean significance across the three cell types.

#### 3.4.2 Functional Enrichment Analysis of Parity Associated Deconvoluted Cell Types

Weighted gene set enrichment meta-analysis was performed on the four parity-responsive cell types using the same four-method approach applied to age and regional effects. Analysis across seven pathway databases identified distinct molecular programs underlying parity-associated cellular composition changes (Figure 5c-e). CA3 pyramidal neurons regulation patterns across datasets were amalgamated through a meta-analysis combining both 356_CA3-do (10x dataset) and 358_CA3-do (Smart-seq dataset) populations revealing shared molecular signatures (Figure 5c). The most significant downregulated pathways included GSK3B and BTRC:CUL1-mediated degradation of NFE2L2 (meta-score = 7.88, NES = - 3.55) and ubiquitin-dependent degradation of Cyclin D (meta-score = 7.86, NES = -3.42), while the strongest upregulated programs were dominated by translation and ribosome biogenesis machinery: eukaryotic translation initiation (meta-score = 6.62, NES = +3.50), SRP-dependent cotranslational protein targeting to membrane (meta-score = 6.51, NES = +3.29), eukaryotic translation elongation (meta-score = 6.49, NES = +3.49), KEGG ribosome (meta-score = 6.17, NES = +3.41), and nonsense-mediated decay (meta-score = 5.86, NES = +3.16).

Astrocytes (376_Astro) displayed predominantly upregulated pathways focused on cellular remodelling and stress response (Figure 5d), with cell-specific signatures including protein-containing complex disassembly (NES = 2.35), protein domain specific binding (NES = 2.21), chemotaxis and taxis responses (NES = 1.89), regulation of neurogenesis (NES = 1.87), and hypoxia response pathways (NES = 1.81).

Sst interneurons (78_Sst HPF) yielded 553 enriched pathways, with 60 reaching Moderate confidence and 22 downregulated pathways at Method-Specific tier (Figure 5e). The strongest upregulated hits centred on broad synaptic and neuronal architecture (synapse meta-score = 4.71, NES = +4.71; plasma membrane bounded cell projection 4.35, +4.35; neuron projection 4.30, +4.30; postsynapse 4.16, +4.16), with cell-specific method-specific signatures including presynaptic membrane (NES = +2.03) and endomembrane system organization (NES = +1.88). Downregulated programs were limited and concentrated on chromatin and RNA-binding processes (chromatin DNA binding NES = -2.84; DNA binding NES = -2.56; RNA metabolic process NES = -2.55).

Cross-cell type pathway analysis revealed shared molecular programs, with 419 pathways enriched across all three cell types focusing on synaptic signaling, cellular respiration, and neurotransmitter transport, while 110 pathways were specifically shared between astrocytes and Sst interneurons, emphasizing cytoskeletal organization and neuroactive ligand signalling. The comprehensive analysis established a hierarchy of molecular disruption: 358_CA3-do (1,557 total enriched pathways, including 345 reaching Moderate confidence or higher) > 376_Astro (570 total / 56 Moderate) ≈ 78_Sst_HPF (553 total / 60 Moderate) > 356_CA3-do (50 total, all method-specific tier), indicating cell type-specific vulnerability to parity with CA3 neurons from the Smart-seq dataset showing the most profound transcriptional remodelling (Figure 5f-i).

#### 3.4.3 Shared Parity Effects in Multiple Affected Cell Types

Comparative pathway analysis identified shared and distinct molecular programs across parity-responsive cell types (Figure 5f-i). Pairwise overlaps revealed selective coordination patterns, with astrocytes and Sst interneurons sharing the most pathways (110 pathways, 6.9%), featuring cytoskeletal organization, MAPK cascade regulation, endocytosis, and neuroactive ligand signalling processes (Figure 5h). CA3 neurons and astrocytes shared 22 pathways (1.4%) emphasizing purine nucleotide biosynthesis, protein kinase regulation, and axonogenesis control (Figure 5f), while CA3 neurons and Sst interneurons showed minimal overlap (11 pathways, 0.7%) focusing on amino acid transport, cell growth regulation, and potassium ion transport (Figure 5g). This pathway distribution pattern indicates that parity triggers predominantly shared molecular responses (26.2% core pathways) with selective cell type partnerships, particularly between astrocytes and Sst interneurons, suggesting coordinated glial-interneuron remodelling as a central feature of parity-induced hippocampal plasticity. Pathway overlap analysis of all 1,600 total enriched pathways revealed extensive coordination between cell types, with 419 pathways (26.2%) shared across all three cell types, emphasizing core parity responses including synaptic signalling, cellular respiration, neurotransmitter transport, and energy metabolism (Figure 5i).

## 4. Discussion

This study performed differential gene expression and cell-type deconvolution analyses on female bulk RNAseq from the dorsal and ventral hippocampus of young and middle-aged rats with differing parity. We identified powerful effects of aging (six cell types), regional heterogeneity (nine cell types), and age × region interactions (two cell types). Parity selectively preserved or increased the proportions of dorsal CA3 pyramidal neurons and somatostatin interneurons, while modestly reducing astrocyte proportions alongside upregulated stress-response and remodelling programs. Our findings add to the growing literature describing long-term signatures of parity in the hippocampus that persist after weaning and into middle age.

### 4.1 Age-related cellular remodelling reveals female-specific vulnerabilities

Four of the six age-significant cell types we identified showed proportional decreases with age: Parvalbumin and vasoactive intestinal polypeptide interneurons (PVALB/VIP; -91%), microglia and perivascular macrophages (386_Micro-PVM; -38%), subiculum neurons (subiculum; -24.8%), and somatostatin interneurons (SST; -22.4%). The concurrent decrease in PVALB interneurons which comprise approximately 24% of GABAergic neurons in CA1 may further compromise network function, and females exhibit twice the vulnerability to PV interneuron loss under stress conditions, potentially contributing to sex differences in age-related cognitive decline and stress-related psychiatric disorders^29^. The 386_Micro-PVM population also declined with age (−38%). This likely reflects the distinction between cellular proportion and activation state, with the homeostatic Micro-PVM signature declining while a distinct aged-microglial phenotype emerges, captured in our analysis as a Microglia class detected almost exclusively in aged samples. Subicular populations showed mixed effects, with the Subiculum class declining (−25%) and Cluster_28_Subi increasing (+17%). SST interneurons, which preferentially target dendritic regions and regulate feedback inhibition, show vulnerability during aging^26^, and inhibition of SST-positive dentate hilar interneurons is sufficient to replicate hallmarks of hippocampal aging, including microglial activation and cognitive deficits^27^. Our findings mirror those of Gramuntell et al. (2021), who demonstrated age-related structural changes in SST interneurons with dendritic spine density alterations appearing as early as 9 months in female mice^28^.

### 4.2 Dorsal-ventral heterogeneity shapes hippocampal cellular architecture

Our deconvolution revealed nine cell types with regional differences along the dorsal-ventral hippocampal axis, showing that this functional specialisation is underpinned by systematic differences in cellular composition that partition into a dorsal compartment dominated by principal projection neurons and oligodendrocytes, and a ventral compartment dominated by inhibitory neurons and microglial-PVM populations. The dorsal hippocampus contained significantly greater proportions of DG/CA1/CA2/CA3_dorsal (p = 6.51 × 10⁻²⁹), CA1/CA2_dorsal (p = 4.69 × 10⁻⁴), and Oligodendrocyte (p = 4.95 × 10⁻¹⁶), whereas the ventral hippocampus contained greater proportions of Sst_IN (p = 4.14 × 10⁻¹⁵), Pvalb/Vip_IN (p = 0.0087), Sncg_IN (p = 0.018), Subiculum (p = 2.57 × 10⁻⁹), Cluster_25_Subi (p = 9.32 × 10⁻¹⁹), and 386_Micro-PVM (p = 3.79 × 10⁻⁴).

The dorsal hippocampus encode more and smaller, more precisely defined place fields than their ventral counterparts^30^, requiring precise excitatory-inhibitory coordination contributed by both SST^31^ and PV interneurons^32^. The ventral enrichment of these inhibitory populations we observe likely reflects the greater inhibitory infrastructure required to gate ventral hippocampal output to limbic targets and support the emotional and affective regulation.

Lodge et al. (2023) provide circuit-level support for this division, showing that within the ventral hippocampus, SST interneurons preferentially regulate pyramidal cells projecting to the nucleus accumbens, directly linking ventral inhibitory infrastructure to limbic-output gating^33^. Oligodendrocytes showed strong dorsal enrichment (p = 4.95 × 10⁻¹⁶), which aligns with single-cell transcriptomic studies showing mature oligodendrocytes are present in unique proportions in each brain area^34^.

The influence of age on the dorsal and ventral hippocampus act through partially distinct cellular mechanics. The 386_Micro-PVM population declined with age in both regions, but more steeply in ventral (−45%) than dorsal (−27%) (age × region p = 0.003), such that the prominent ventral enrichment of microglial-PVM in young animals was largely lost by middle age. Oligodendrocytes showed the opposite regional aging pattern, increasing modestly in dorsal (+7%) and decreasing in ventral (−9%) (p = 0.013), accentuating their dorsal enrichment with age^35^. The regional heterogeneity of these aging trajectories is consistent with spatial transcriptomic evidence that age-associated neuroinflammatory and myelin-related changes concentrate in regions of high myelinated fibre density^36^. Functional enrichment of the 386_Micro-PVM age × region interaction identified high confidence pathways in signalling receptor activity and molecular transducer activity, suggesting that the regional aging trajectory involves active microglial signalling reorganisation rather than passive cellular loss.

### 4.3 Parity fundamentally reorganizes hippocampal cellular composition

Although age and region dominated variance in hippocampal cellular composition, parity emerged as an independent modulator of three specific populations: dorsal CA3 pyramidal neurons, SST interneurons, and astrocytes.

#### 4.3.1 CA3 pyramidal neurons

Dorsal CA3 pyramidal neurons showed small but consistent parity-associated increases across both reference datasets (+4.4% in 358_CA3-do, +8.7% in 356_CA3-do). Pregnancy and the early postpartum are associated with acute structural remodelling of CA3, including reduced dendritic branching and length^37,38^, whereas our data indicate preserved or modestly elevated CA3 proportions at the postpartum and middle age timepoints examined here. CA3 pyramidal neurons are critical for pattern completion and the retrieval of memory^39^, and parity is associated with improved memory persisting long past the reproductive period^40–43^. Pathway analysis revealed downregulation of protein degradation programs, specifically GSK3B/BTRC:CUL1-mediated degradation of NFE2L2 and ubiquitin-dependent Cyclin D degradation, alongside upregulation of translation and ribosome biogenesis machinery, including eukaryotic translation initiation, ribosome assembly, and cotranslational protein targeting. Together this bidirectional pattern suggests that parity triggers a coordinated shift in CA3 proteostatic balance, reducing protein turnover while increasing protein synthesis capacity in the preserved population. In both rodents and humans, pregnancy and the postpartum is associated with dramatic changes in hippocampal volume^44^ that persist for at least 6 years following birth^45,46^, including specific volume reductions in CA2/CA3 subfields that followed a distinct trajectory from CA1 changes^47^. Along with hippocampal volume decreased, striking reductions are seen in hippocampal neurogenesis across the postpartum into the later postpartum that are reversed and increased into middle-age, at least consistent with the downregulation of expression of genes involved in the degradation of cyclin D we observed here in dorsal CA3^9,12^. Our cell-type-level data add resolution to these findings, indicating that the persistent subfield-level volume changes in humans may coexist with preservation of CA3 cell number at the postpartum and middle-age timepoints examined.

#### 4.3.2 SST interneurons

Hippocampal SST interneurons (78_Sst HPF) showed a consistent parity-associated increase in proportion (+8.1%), a directional pattern opposite to the marked age-related SST decline (− 22%) observed in our own data. SST interneurons regulate dendritic integration and feedback inhibition through preferential targeting of pyramidal cell dendrites^31^, and their loss or dysfunction has been causally linked to age-associated cognitive decline. Lyu et al. (2023) demonstrated that selective chemogenetic inhibition of SST-positive dentate hilar interneurons alone in male mice was sufficient to replicate hallmarks of hippocampal aging, including microglial activation and spatial memory impairments^27^. This suggests that the parity-associated SST increase we observe would be predicted to support rather than degrade hippocampal inhibitory and cognitive function, consistent with the established behavioural signature of preserved cognition in parous rodents and humans through middle age^42,59,60^.

The molecular mechanisms underlying parity-associated SST changes likely involve pregnancy-driven reorganisation of GABAergic signalling. During pregnancy, the GABA system undergoes extensive modification, largely driven by allopregnanolone fluctuations^48^ producing a neurosteroid withdrawal state resulting in reduced δ-GABAAR expression on hippocampal interneurons during pregnancy which reverts within 48 hours postpartum^49,50^. Yet, our data, collected at 30 days and 6 months postpartum, suggest that some component of this GABAergic reorganisation persists well beyond the acute peripartum period. Our pathway analysis supports this interpretation: upregulated pathways in SST interneurons featured broad synaptic and neuronal architecture, including presynaptic membrane organisation and endomembrane system organisation, while the limited downregulated signal concentrated on chromatin and RNA-binding processes. This pattern is consistent with a model in which parity induces lasting expansion of the SST interneuron population accompanied by reorganisation of synaptic architecture, potentially reflecting adaptive adjustment of inhibitory tone following the dramatic GABAergic modulation of pregnancy.

#### 4.3.3 Astrocytes

Astrocyte proportions showed a small but consistent parity-associated decrease (376_Astro, - 1.8%), but pathway analysis revealed exclusively upregulated programs focused on cellular remodelling and stress response. This combination of fewer cells in proportion, but with elevated activity profiles points to a recalibration of the astrocytic compartment toward a leaner, more transcriptionally active state rather than uniform expansion or loss. The specific upregulation of chemotaxis and hypoxia response pathways connects to the established role of astrocytes as metabolic and homeostatic sensors, while pathway hits in protein-containing complex disassembly and protein domain specific binding suggest active proteostatic remodelling.

This pattern aligns with a dynamic astrocytic trajectory across the peripartum and postpartum periods. Astrocyte density is reduced during pregnancy^51^, and our data show that this depletion persists at modest magnitude into the late postpartum and middle-age timepoints we examined, consistent with the down-arm of the U-shaped grey-matter trajectory documented across pregnancy and recovery in humans^52^. Convergent evidence from our prior work shows that previous parity prevents age-related decreases in neural stem cells in the dentate gyrus and maintains higher synaptic protein (PSD95) levels, while parous females maintain enhanced cognitive performance compared to nulliparous controls in normal aging^22^. Recent population-scale lab-test data add further context: Bar et al. (2025) tracked 76 lab tests across more than 300,000 pregnancies and found that systemic inflammation markers remain elevated above preconception baseline well beyond 80 weeks postpartum, with about half of tests requiring 3 months to a year to return to baseline^53^. The persistent astrocytic remodelling we observe at 30 days and 6+ months postpartum sits squarely within this prolonged systemic recovery window, suggesting that hippocampal cellular remodelling is one component of a broader, slow-resolving postpartum physiological state. Whether the leaner, more transcriptionally active astrocyte profile we observe contributes to lasting resilience against age-related cognitive decline or reflects a continuing compensatory response to pregnancy-associated cellular stress remains an open question.

### 4.4 Limitations

Several limitations should be considered when interpreting these findings. First, deconvolution-derived proportion changes are inferred estimates. Whilst our age-related interneuron losses show convergent directionality with the established literature on selective interneuron vulnerability in aging^26,28,29^, convergent evidence is not equivalent to direct validation. Second, our deconvolution used mouse single-cell references to deconvolve rat bulk RNA-sequencing, relying on ortholog mapping between species. Cross-species deconvolution from mouse references to rat bulk RNA-seq is established practice and performs comparably to within-species methods via ortholog mapping^54,55^, but species-specific cell states or genes without clear orthologs may introduce noise. Third, a fundamental limitation of bulk RNA-seq deconvolution is the inability to distinguish genuine changes in cell number from changes in cell-type-specific gene expression magnitude. If, for instance, CA3 neurons upregulate marker genes following parity without undergoing cellular proliferation, this would appear as an increased proportion in deconvolution output. Compounding this, neurons contain as much as 10 times more RNA per cell than glial cells, which systematically biases proportion estimates toward neuronal populations^56^. Shifts in RNA content per cell with parity or aging could therefore manifest as apparent proportional changes without corresponding changes in absolute cell number. To address this our comparisons are relative (between conditions within the same tissue type) rather than absolute, which partially mitigates this concern. Lastly, our data generation occurred in two batches per age grouping. However, many of our identified results match the literature for known age effects as discussed above, and age was used as a covariate in our parity-specific DESeq2 analysis, which also acts to remove this potential batch effect.

### 4.5 Implications and future directions

The intersection of aging, regional specialization, and parity creates a complex landscape of hippocampal plasticity unique to females^57^. Our identification of cell-type-specific parity effects operating independently of age and region establishes parity as a distinct biological variable in hippocampal remodelling, rather than a modifier of existing aging or regional programs. These findings contribute to a growing body of evidence that reproductive experience leaves lasting cellular signatures in the brain. The specific populations affected (CA3 pyramidal neurons, SST interneurons, and astrocytes) are each independently implicated in cognitive aging and neuropsychiatric vulnerability. The parity-associated SST increase we observe runs directly counter to the marked age-related SST decline detected in our own data positioning parity as a candidate cellular contributor to the cognitive resilience documented in middle-aged parous animals. The fact that parity targets these populations raises important questions about how reproductive history intersects with later-life brain health, particularly given epidemiological evidence linking parity number to dementia risk^58^ and neuroimaging data showing both acute grey matter reductions and longer-term neuroprotective signatures in parous women^11,59^. Finally, the convergence between our rodent deconvolution and human subfield-level neuroimaging motivates application of similar deconvolution approaches to existing human postpartum bulk RNA-seq datasets from brain banks, which could identify conserved cellular signatures of reproductive experience across species. Understanding how reproductive experience shapes the brain’s cellular architecture throughout the female lifespan represents a critical frontier in neuroscience with profound implications for women’s health.

## Statements and Declarations

None.

## Author Contribution

**Andrew J McGovern:** Conceptualization, Software, Formal analysis, Writing - Original Draft, Visualization. **Paula Duarte-Guterman:** Conceptualization, Methodology, Resources, Writing - Review & Editing. **Liisa AM Galea:** Conceptualization, Resources, Writing - Review & Editing, Supervision.

## Ethics and Consent to Participate declarations

All protocols were approved by the Institutional Animal Care Committee at the University of British Columbia and conformed to the guidelines set out by the Canadian Council on Animal Care.

## Data Availability

Raw and processed bulk RNA-sequencing data generated for this study have been deposited in the NCBI Gene Expression Omnibus (GEO) under accession number GSE329776. All analysis code is available at https://github.com/AJMcGovernLab/ReproductiveExperienceAndAgeDeconvolution.

## Acknowledgments

We wish to thank Dr Stephanie E. Lieblich for contributions to the development of the parity dataset.

