## Supplement for "Cell Type-Specific Remodelling of the Rat Hippocampus by Parity and Age"

Section numbering mirrors the manuscript (S2.X ↔ §2.X).

**GEO accession:** [GSE329776](https://www.ncbi.nlm.nih.gov/geo/query/acc.cgi?acc=GSE329776)

**Code repository:** <https://github.com/AJMcGovernLab/ReproductiveExperienceAndAgeDeconvolution>

#### Software and package versions

| Component | Version | Purpose |
| --- | --- | --- |
| BCL2FASTQ | 2.20.0.422 | Demultiplexing of Illumina BCL output |
| Cutadapt | (latest at time of analysis) | Adapter and quality trimming |
| FastQC | 0.11.8 | Pre- and post-trim sequence QC |
| STAR | 2.6.1a_08-27 | Read alignment to *Rnor_6.0* (Ensembl 84) |
| HTSeq-count | 0.11.0 | Per-gene read quantification |
| SCDC | 0.0.0.9000 | Cell-type deconvolution |
| DESeq2 | 1.40.0 | Variance-stabilizing transform; binary parity DE |
| clusterProfiler | 4.8.0 | GSEA against GO / KEGG / Reactome / MSigDB |
| randomForest | 4.7-1.1 | Parity-classification feature selection |
| tidyverse | 2.0.0 | Data wrangling |
| ggplot2 | 3.4.2 | Visualization |

Original analysis under **R 4.3.0**; the public reproduction pipeline is verified under **R 4.5.3**. Random seeds, environment manifests, and all driver scripts are checked in to the project repository.

#### S2.3 Reference dataset cell counts and performance metrics

Per-reference cell counts in each sex configuration:

| Reference | Platform | Female cells | Male cells | Mixed cells |
| --- | --- | --- | --- | --- |
| Allen 10X Genomics whole-brain (2020) | 10X Chromium | 22,307 | 59,108 | 81,964 |
| Yao hippocampus 10X | 10X Chromium | 26,314 | 62,272 | 88,674 |
| Mouse-brain Smart-seq2 (2019, NeMO dat-iye7gkp) | Smart-seq2 | 3,693 | 1,849 | 5,869 |

Configurations were compared (Figure 1) on four size-normalized metrics:

- Shannon entropy per 1,000 reference cells: −Σ(pᵢ × log₂(pᵢ)) / 1,000, where pᵢ is the proportion of cell type *i*.
- Relative cell-type detection rate: proportion of maximum detectable cell types identified in each configuration.
- Modified Simpson's diversity index: standard Simpson's index normalized by log₁₀(n_ref_cells + 1) to control for reference size.
- Detection efficiency: cell types detected per 1,000 reference cells.

Female-matched configurations consistently outperformed mixed-sex and male-only configurations on the 10X-based references; all deconvolution used the female-only configuration.

#### S2.4 Cell-type harmonization detail

The harmonization computed pairwise Pearson correlations between all 349 cell-type signatures (extracted from the SCDC basis matrices), then performed hierarchical clustering with average linkage on the distance matrix 1 − r at a height cutoff of 0.2. Per-reference reduction:

| Reference | Originals | Harmonized |
| --- | --- | --- |
| mouse10x_2020 | 169 | 16 |
| mouse_smartseq_2019 | 125 | 10 |
| yao_hippo_10x | 55 | 1 |
| Total (sum across datasets) | 349 | 27 |

The 27 entries are the per-dataset-summed harmonized categories (Figure 2A).

Two mapping files are used at different stages:

- cell_type_mapping_table.csv for first-pass harmonization (Figure 2A / 2.4 counts).
- refined_cell_type_mapping.csv for second-pass refinement (additional h = 0.1 sub-clustering + biological renaming), used as the row identity for the harmonized three-way ANOVA.

Within- vs between-cluster correlation statistics (reported in 3.2) are in correlation_summary_stats.csv.

#### S2.5 SCDC deconvolution parameters

- Subject parameter: animal ID, to account for biological replicates and improve estimation stability.
- Ensemble weighting: equal weights across the three reference datasets; sensitivity analyses with alternative weighting schemes produced concordant proportions (Pearson r > 0.95).
- Convergence and reconstruction errors were verified on every sample.
- Technical-replicate correlation consistently exceeded r = 0.85.

Proportions estimated from harmonized vs. non-harmonized annotations were compared to assessing the impact of standardization on biological inference.

#### S2.6 Directional inference for cell-type proportions

For directional inference of significant main effects, marginal log2 fold change estimates were computed as the unweighted mean of within-stratum log2 fold changes across the 4 (age × region) or (age × parity) cells of the design. This conditioned estimator matches the directional component of the three-way ANOVA p-value, in contrast to a simple unconditioned pooled contrast (e.g. parous vs nulliparous across all samples), which is biased by variance attributable to the other two factors when the design is unbalanced. Per-stratum signs accompany each directional log2 fold change in the results as a non-parametric robustness check. For cross-dataset consensus: a finding was high-confidence when it showed the same direction and p < 0.05 across ≥ 2 reference datasets; cross-dataset combination used Fisher's combined probability test with adjustment for between-dataset correlation.

#### S2.7 Differential gene expression

##### S2.7.1 Type III sums of squares

The gene-level three-way ANOVA (expression ~ Age × Region × Parity, aov()) used Type III sums of squares, justified by the unbalanced factorial design; Benjamini-Hochberg FDR < 0.05 was applied within each of the seven model terms.

##### S2.7.2 Enrichment weighting input (DESeq2)

For functional enrichment, weighted gene scores combined effect magnitude and statistical confidence:

weight = 0.5 × |log2(fold-change)| + 0.5 × (−log10(adjusted p-value))

##### S2.7.3 PCA driver-gene identification

PCA was performed on the variance-stabilized (DESeq2 VST) expression matrix; the top 10 PCs explained > 60% of total variance. Each PC was tested for factor associations with seven nested linear models (main effects and all interaction combinations). PC5, PC6, and PC8 were significantly associated with binary parity (Parity_Binary main-effect model: p = 0.048, 0.026, 0.013; R² = 0.066, 0.083, 0.101).

For parity-associated PCs, driver genes were identified by ranking genes by absolute loading and quantifying each gene's contribution as the reduction in model R² when its influence on the PC was removed: the score reduced = pc_scores − (gene_loading × gene_expression) was substituted for the original PC scores in the parity-binary model, and the R² loss (r2_loss) and parity-coefficient p-value inflation (log_p_degradation) were combined into a contribution score log_p_degradation + 100 × r2_loss; genes above the elbow of that distribution were retained as drivers:

| PC | Driver genes |
| --- | --- |
| PC5 | 63 |
| PC6 | 128 |
| PC8 | 168 |

##### S2.7.4 Random-forest parameters and reproducibility

The multi-class parity label (nulliparous, primiparous, biparous; 3 classes) was the response variable instead of the binary grouping used elsewhere to allow the classifier to leverage primiparous–biparous differences which out-performed the binary model.

- 1,000 decision trees (ntree = 1000); bootstrap sampling per tree; mtry = floor(√(n_features)) = **111**; set.seed(12345).
- Mean Decrease in Accuracy (MDA) = drop in classification accuracy when each variable is permuted; Mean Decrease in Gini (MDG) = total node-impurity decrease averaged across trees.
- The number of parity-predictive genes (23) was determined from the cross-validation classification accuracy across candidate gene-set sizes, which peaked at 23 genes (5-fold cross-validation, 3 repeats; Supplementary Figure 5). The top 23 genes by MDA were retained; saved-model MDA range 1.83–3.04 (top ENSRNOG00000053712 = 3.036; bottom Vrk3 = 1.832).

Note: The trained model objects are the canonical artefacts for the top-23 list (loading them reproduces it exactly on any platform). The pipeline (07_suppfig234_parity_genes/02_random_forest.R) defaults to loading the saved models; REPRO_RF_REFIT=TRUE retrains from scratch.

#### S2.8 Functional enrichment: Weighting, GSEA, and Meta-Analysis

##### S2.8.1 Weighting schemes and databases

Four complementary weighting schemes integrated evidence from the DESeq2, PCA, and random-forest analyses:

| Scheme | Composition |
| --- | --- |
| Association-weighted | 100% gene–cell-type association score |
| Expression-weighted | 60% association + 40% \|log₂(fold-change)\| |
| Statistical-weighted | 60% association + 40% −log₁₀(FDR) |
| Composite-weighted | 50% association + 25% \|log₂(fold-change)\| + 25% −log₁₀(FDR) |

Seven pathway database collections were queried: GO:BP, GO:MF, GO:CC, KEGG, Reactome, and two MSigDB collections (Hallmark and Curated). For each cell type, weighting scheme, and percentage-rank cutoff (top 5% / 10% / 25% by signed weighted score, contributing meta-analysis weights 0.50 / 0.40 / 0.10), the ranked gene list was submitted to GSEA against each database.

##### S2.8.2 GSEA implementation

GSEA used clusterProfiler v4.8.0: gseGO() for the three GO ontologies, gseKEGG() for KEGG, gsePathway() (ReactomePA) for Reactome, and GSEA() for the MSigDB collections.

- Minimum gene-set size 5; maximum 500.
- Default gene-set permutation (clusterProfiler fgsea backend); Benjamini-Hochberg adjustment.
- Pathway magnitude = normalised enrichment score (NES), comparable across gene-set sizes; sign encodes the leading-edge position in the ranked input.
- Directional interpretation: because inputs are ranked by signed weighted score (positive = parity-upregulated), positive NES = upregulated, negative NES = downregulated in the relevant cell type.

##### S2.8.3 Meta-score and confidence tiers

For each pathway:

meta-score = consensus × evidence × max(0, NES_consistency)

- consensus = proportion of methods returning the pathway with FDR < 0.05.
- evidence = weighted mean −log₁₀(method-level FDR), method weights {association 0.30, expression 0.25, statistical 0.25, composite 0.20}.
- NES consistency = 1 − (SD / (|mean| + ε)) of per-method NES, clamped at ≥ 0.
- Method p-values combined via Fisher's combined probability test (pchisq(−2 × Σ log(p), df = 2·n_methods)); the resulting meta-FDR was used for filtering.

| Tier | Method count | Meta-score |
| --- | --- | --- |
| Ultra-high | All 4 | > 7 |
| High | ≥ 3 | > 5 |
| Moderate | ≥ 2 | > 3 |
| Method-specific | 1 | — |

A pathway was classified as a cell-type-specific if enriched in ≤ 2 cell types with a meta-score difference > 2 between enriched and non-enriched cell types.

#### Cross-reference index

| Manuscript Section | Driver script directory |
| --- | --- |
| 2.3 | scripts/01_fig1_reference_sex/ |
| 2.4 | scripts/02_fig2_harmonization/ |
| 2.5 | scripts/01_fig1_reference_sex/01_run_deconvolution.R (SCDC; outputs staged in checkpoints/scdc_deconvolution/) |
| 2.6 | scripts/03_fig3_harmonized_anova/ (harmonized) and scripts/06_fig5ab_parity_proportions/ (per-dataset) |
| 2.7.1 | scripts/04_suppfig1_gene_anova/ |
| 2.7.2 | scripts/07_suppfig234_parity_genes/01_deseq2_binary_parity.R |
| 2.7.3 | scripts/06b_pca_preprocessing/ and scripts/07_suppfig234_parity_genes/03_pc_driver_analysis.R |
| 2.7.4 | scripts/06c_rf_preprocessing/ and scripts/07_suppfig234_parity_genes/02_random_forest.R |
| 2.8 | scripts/05_fig4_age_region_enrichment/ (age/region) and scripts/08_fig5ce_parity_enrichment/ (parity); overlaps in scripts/09_fig5fi_pathway_overlaps/ |

**End-to-end reproduction:** Rscript scripts/run_all_robust.R from the Repository root regenerates files in outputs/ from the staged data and checkpoints. SCDC deconvolution (§2.5) is checkpointed because SCDC is not on CRAN (remotes::install_github("meichendong/SCDC")); its outputs are pre-staged in checkpoints/scdc_deconvolution/. See manuscript/DATA_PROVENANCE.md for per-file provenance.

Supplementary Figures

Cell Type-Specific Remodelling of the Rat Hippocampus by Parity and Age

McGovern AJ, Duarte-Guterman P, Galea LAM

### Supplementary Figures:

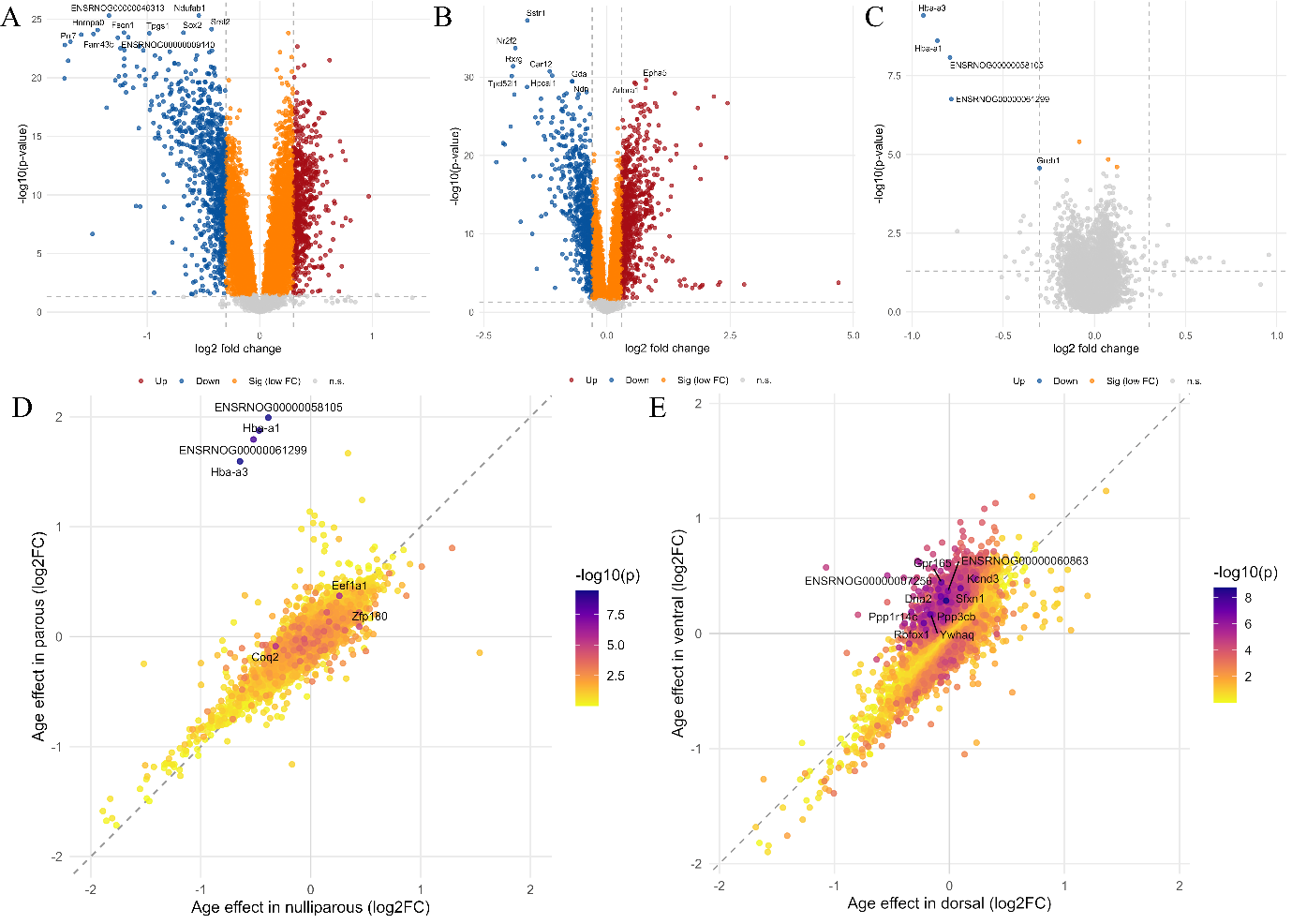

**Supplementary Figure 1:** Three-way ANOVA differential genes expression analysis found pronounced effects of (A) age and (B) region, but the effect of parity itself was limited with 8 genes achieving statistical significance (FDR<0.05). There was a small (D) parity×age effect (7 genes), and a pronounced (E) age×region effect, and but no significant parity×region effect, or three-way interactions.

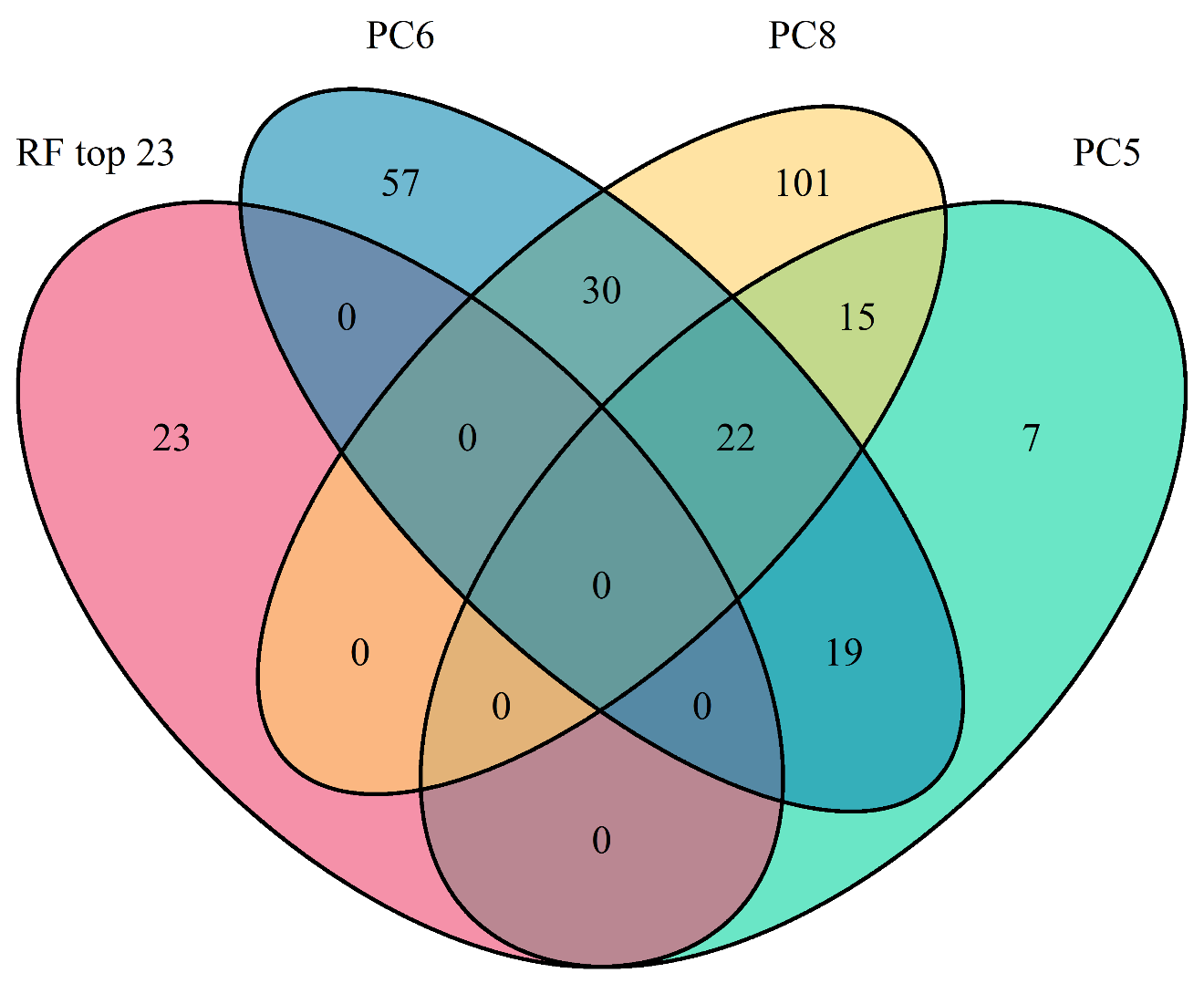

**Supplementary Figure 2:** Overlap Between Random Forest and PCA Parity Genes. Four-way venn diagram of parity associated genes identified by principal component factor associations (PC5, PC6, and PC8) with the genes identified as predictive of parity by random forest modelling.

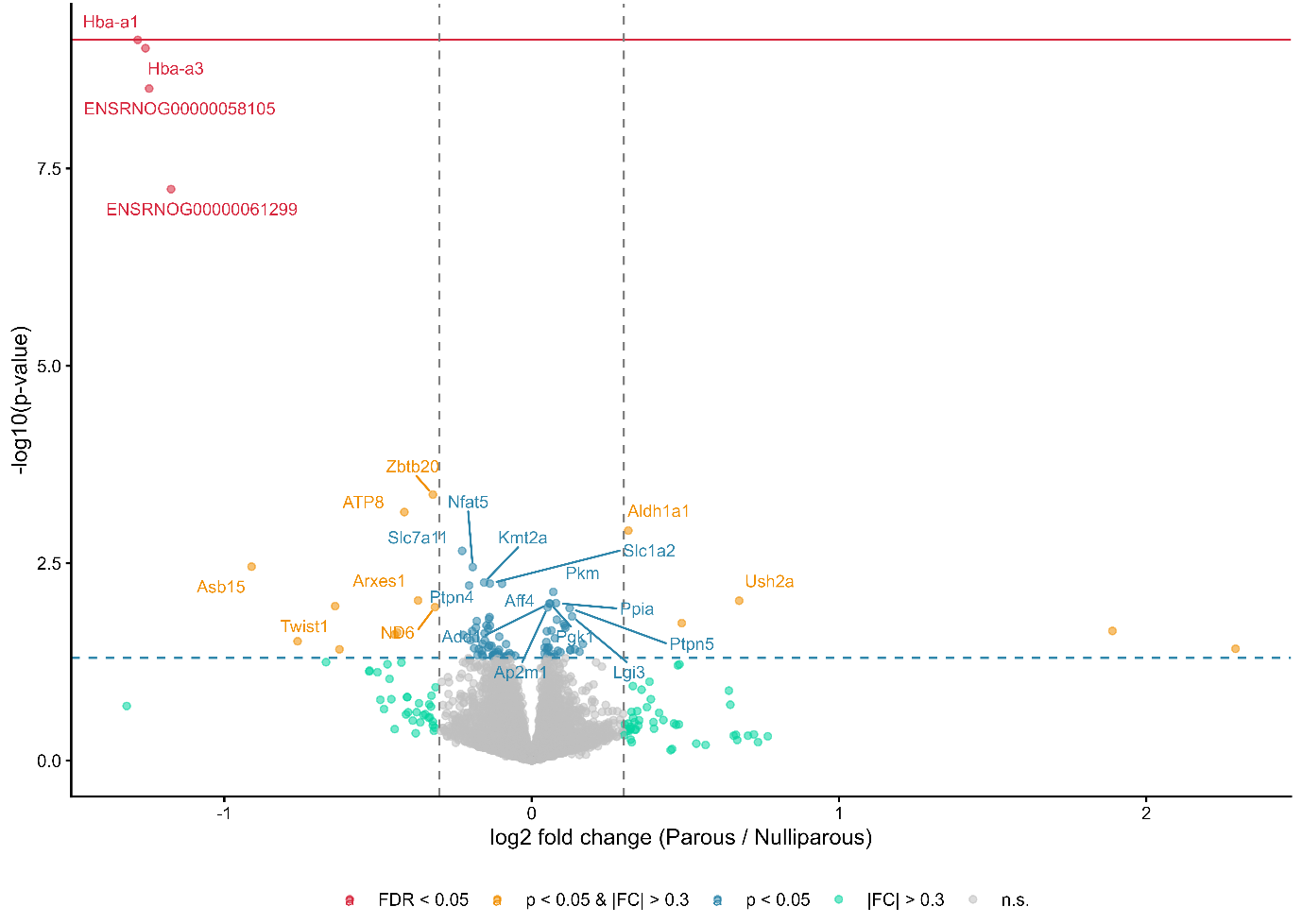

**Supplementary Figure 3:** Volcano of Binary Parity DESeq2. Volcano plot showing differential gene expression between parous (biparous and primiparous combined) and nulliparous rats across the hippocampus. Analysis performed using DESeq2 with age as a covariate (design: ~ age + parity). X-axis represents log2 fold change (Parous/Nulliparous); y-axis shows -log10(p-value). Points colored by significance: red (FDR < 0.05), purple (FDR < 0.2), orange (p < 0.05 and |FC| > 0.3), blue (p < 0.05), green (|FC| > 0.3), gray (not significant). Horizontal lines indicate p = 0.05 (dashed blue) and FDR = 0.05 threshold (solid red); vertical dashed lines mark fold change = ±0.3. Analysis included genes with mean count ≥1, variance >0, detected in ≥6 of 60 samples (≥10%) from the filtered expression matrix.

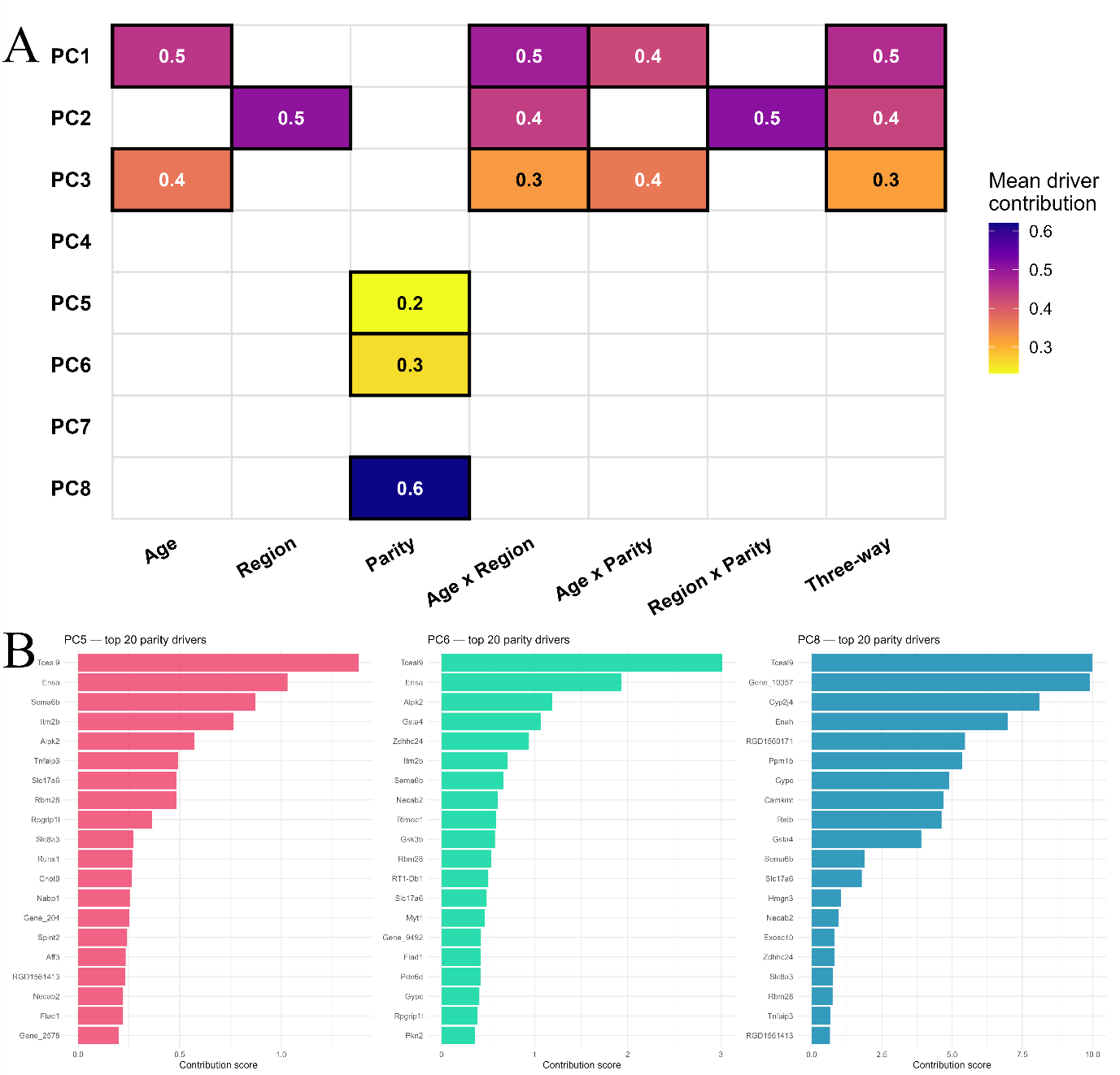

**Supplementary Figure 4:** (A) Principal component factor analysis identifies pc5, 6, and 8 to be significantly influenced by parity. (B) Permutation testing identified genes contributing to statistical significance of parity in these components.

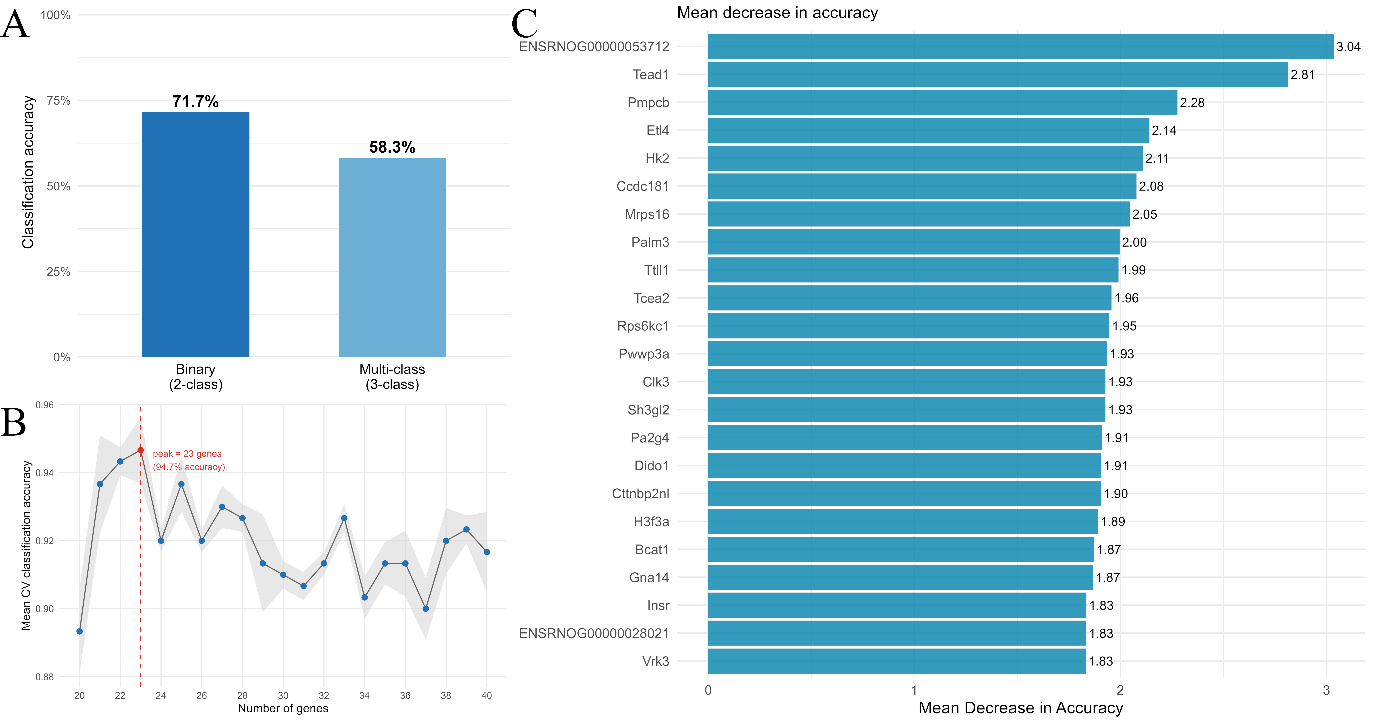

**Supplementary Figure 5:** (A) A binary random forest machine learning model achieves 71.7% accuracy. (B) An elbow curve on the mean cross-validation accuracy identifies (C) 23 genes as contributing most efficiently for the model.
